# Increasing Ciliary ARL13B Expression Drives Active and Inhibitor-Resistant SMO and GLI into Glioma Primary Cilia

**DOI:** 10.1101/2022.11.28.518234

**Authors:** Ping Shi, Jia Tian, Julianne C. Mallinger, Loic P. Deleyrolle, Jeremy C. McIntyre, Tamara Caspary, Joshua J Breunig, Matthew R. Sarkisian

## Abstract

ADP-ribosylation factor-like protein 13B (ARL13B), a regulatory GTPase and guanine exchange factor (GEF) enriches in primary cilia and promotes tumorigenesis in part by regulating Smoothened (SMO), GLI, and Sonic hedgehog (SHH) signaling. Gliomas with increased *ARL13B, SMO* and *GLI2* expression are more aggressive but the relationship to cilia is unclear. Previous studies showed increasing ARL13B in glioblastoma cells promoted ciliary SMO accumulation, independent of exogenous SHH addition. Here we show SMO accumulation is due to increased ciliary, but not extraciliary ARL13B. Increasing ARL13B expression promotes the accumulation of both activated SMO and GLI2 in glioma cilia, but not in NIH3T3 fibroblast cilia. ARL13B-driven increases in ciliary SMO and GLI2 are resistant to SMO inhibitors, GDC-0449 and cyclopamine. Finally, temozolomide chemotherapy which increases ARL13B expression in glioma, stimulates SMO and GLI2 into glioma cilia, but not fibroblast cilia. Collectively, our data suggest factors that elevate ARL13B may drive drug-resistant SMO and GLI into cilia. This suggests the ARL13B-associated mechanism that leads to ciliary SMO/GLI recruitment may promote treatment resistance in glioma.

## Introduction

Glioblastoma (GBM) is the most lethal form of primary brain tumors in adults, presumably originating from a transformed glial cell lineage (Omuro and DeAngelis, 2013). GBMs also evolve from lower grade gliomas(Aiman and Rayi, 2022; Jooma et al., 2019; Osborn et al., 2022). GBMs inevitably recur and average patient survival from diagnosis averages 15-18 months despite standard of care therapies (surgery, temozolomide (TMZ) chemotherapy, and irradiation) (Stupp et al., 2005; Tykocki and Eltayeb, 2018). Tumors have complex modes of signaling that allow them to escape therapy and promote malignancy, including changes in gene expression patterns and signaling pathway alterations. How cellular structures contribute to these signaling pathways remains unclear. The primary cilium is a relatively understudied organelle in glioma, and a major signaling structure of many cell types, normal and malignant (Alvarez-Satta and Matheu, 2018). Cilia concentrate several signaling pathways - the best characterized being Sonic hedgehog (SHH)(Wheway et al., 2018). SHH is one of three types of vertebrate hedgehog (Hh) ligands that stimulate cell proliferation, survival and migration of normal and cancer cells (Han and Alvarez-Buylla, 2010; Liu et al., 2018). While the SHH pathway appears overactive in a significant fraction of GBMs (Bar et al., 2007a; Bar et al., 2007b; Clement et al., 2007; Gruber Filbin et al., 2013; Xu et al., 2008), little research has been done to understand the relationship of SHH and cilia in GBMs.

GBM tumors and derived cell lines from patients can display up to ~30% ciliated cells at any given time (Goranci-Buzhala et al., 2021; Hoang-Minh et al., 2016b; Hoang-Minh et al., 2018; Lee et al., 2022; Sarkisian et al., 2014; Wei et al., 2022). Like other cell types, glioma cilia enrich ARL13B (Hoang-Minh et al., 2018; Sarkisian et al., 2014), a regulatory GTPase and GEF localized to the ciliary membrane required for ciliary structural maintenance (Caspary et al., 2007). Recent studies suggest ARL13B is critical for tumor growth in vitro and in vivo, as well as angiogenesis within GBM (Chen et al., 2022; Hoang-Minh et al., 2018; Lee et al., 2022; Shi et al., 2021; Shireman et al., 2021)Shireman 2021. For example, mice carrying intracranial GBM tumors expressing dominant-negative KIF3A lacked ARL13B+ cilia and displayed prolonged survival (Hoang-Minh et al., 2016b). Knockdown of ARL13B within all GBM subtypes slowed intracranial tumor growth in vivo (Shireman et al., 2021). More recently, signaling thru or at ARL13B+ cilia appears critical for cells to maintain a proliferative or stem like state through engagement of the SHH pathway. For example, histone deacetylase 6 (HDAC6) inhibition can trigger glioma stem cell differentiation by suppressing SHH signaling (Yang et al., 2018). However this phenomenon is not observed in glioma cells depleted with ARL13B (Shi et al., 2021). Similarly, the superenhancer KLHDC8a stimulates glioma stem cell growth by promoting ciliogenesis and SHH pathway activation. Inhibition of ARL13B with shRNA or ciliobrevin, slowed tumor growth, reduced stem cell markers and SHH pathway activation (Lee et al., 2022). Thus ARL13B appears to play a key role in regulating GBM growth and maintaining glioma stem cells.

ARL13B in GBM cilia may also be critical for mediating resistance to chemotherapy. ARL13B induction by TMZ appears to be highly involved in the chemoresistance pathway in GBM(Shireman et al., 2021). TMZ-stimulated expression of ARL13B epigenetically through EZH2, is associated with increased numbers and length of ARL13B+ cilia. This group found that knocking down ARL13B in patient xenografts increased sensitivity to TMZ in vivo (Shireman et al., 2021). These findings are consistent with earlier studies in which depletion of ARL13B+ cilia via CRISPR/Cas9 ablation of *KIF3A* or *PCM1* resulted in increased TMZ sensitivity (Hoang-Minh et al., 2016a). Recently, tumor treating fields (TTFields) ablation of ARL13B+ cilia on GBM cells not only blocked TMZ-induced ciliogenesis but sensitizes GBM cells to TMZ(Shi et al., 2022). Altogether, these results support a role for ARL13B in glioma growth, and environmental factors such as standard of care chemotherapy may drive unwanted changes in ARL13B that could enhance treatment resistance.

ARL13B is a mediator of SHH signaling in the cilium, as loss or mutation of ARL13B alters the enrichment of SHH signaling components in cilia along with downstream transcriptional signaling (Larkins et al., 2011; Mariani et al., 2016). Canonically, SHH binds its receptor, Patched (i.e. *PTCH1* and *PTCH2* genes), on the ciliary membrane and the complex is trafficked to the cell. The G protein-coupled receptor Smoothened (SMO) is then visibly enriched in cilia where it is activated. SMO activation drives the activation of GLI transcription factors, notably GLI1 and GLI2, (Goetz and Anderson, 2010; Goetz et al., 2009), resulting in transcriptional changes in the nucleus (Kim et al., 2009; Santos and Reiter, 2014). This pathway is inhibited by blocking processes leading to the activation of SMO including mutations altering cilia function (Bangs and Anderson, 2017). However, knockout of *Arl13b* in a mouse model results in constitutive SMO ciliary enrichment and SHH ligand-independent pathway activation (Caspary et al., 2007; Larkins et al., 2011; Lu et al., 2015). Further, a single point mutation, V358A, prevents ARL13B from enriching in cilia (Gigante et al., 2020; Mariani et al., 2016), but preserves its GTPase and GEF activity (Gigante et al., 2020). *Arl13b*^*V358A/V358A*^ mice are viable, and surprisingly display normal SHH signaling and ciliary SMO enrichment (Gigante et al., 2020). Thus, how these findings relate to glioma cells and whether the relationship between ARL13B and SMO is SHH-dependent remains unclear.

Hyperactive forms of SMO signaling and ciliary localization are oncogenic (Han and Alvarez-Buylla, 2010), and thus SMO signaling must be tightly controlled. The Cancer Genome Atlas (TCGA) data sets indicate gliomas with high *ARL13B* and *SMO* mRNA expression correlate with shorter overall patient survival (Hoang-Minh et al., 2018; Shireman et al., 2021). Whether high *ARL13B* and *SMO* expression are coincidental or related is not clear. Our previous findings implicated a relationship between ARL13B and SMO in glioma cells that may, in part, be independent of SHH. In one of our GBM patient-derived cell lines (L0) expressing *Arl13b:GFP* transgenes (Hoang-Minh et al., 2018), we observed that glioma cells displayed elongated cilia, accumulation of SMO at their ciliary tips, and excision of their distal tips. This SMO accumulation in *Arl13b:GFP* transgene-expressing cells occurred in the absence of SHH stimulation and was as robust as SHH-stimulated parental cells (Hoang-Minh et al., 2018). However, we do not know whether the ARL13B-mediated changes in glioma cilia length or SMO distribution are attributable to ARL13B’s ciliary or cellular role. Here, we used additional cell lines and known mutations that affect ARL13B subcellular localization and GEF activity. Our goal was to try and distinguish whether ARL13B inside or outside of glioma cilia drives the ciliary changes in SMO, and whether ARL13B drives increases in activated ciliary SMO. Finally, from a clinical perspective, we asked whether ARL13B-driven increases in ciliary SMO are resistant to SMO inhibitors and, whether TMZ-stimulated ARL13B expression enhances ciliary localization of SMO or GLI2 in glioma cells.

## Results

### ARL13B^+^ cilia are present in GBM biopsies and recently derived patient cell lines

Previous studies of widely used GBM cell lines (e.g. U-lines), found that primary cilia are typically absent, structurally impaired or rarely elongated (Moser et al., 2009). We immunostained these cells to examine the presence of ARL13B^+^ cilia. Our experiments support previous findings, and extend them to other widely used glioma lines (A172, F98) (**Suppl. Fig. 1**). However, using biopsy material, recently derived low- or high-grade human glioma cell lines, and high-grade mouse glioma cells (KR158) (Gursel et al., 2011), we, and others, find ARL13B^+^ cilia are readily identifiable (Hoang-Minh et al., 2018; Sarkisian et al., 2014; Shireman et al., 2021; Zalenski et al., 2020) (**Suppl. Fig. 1**). Thus, while previous results may represent some types of gliomas, there may be an association of ARL13B and the cilium with other aggressive gliomas.

### Increasing *ARL13B* expression correlates with increasing *SMO* and *GLI2* and more aggressive glioma

We first examined the relationship of *ARL13B* expression and another key ciliogenesis gene, *IFT88*, with the expression of other SHH pathway genes (*SMO, GLI2* and *SHH*) in low grade (LGG) and high grade (GBM) using TCGA datasets. Strikingly, we find a significant positive correlation between ARL13B and SMO, as well as between *ARL13B* and *GLI2* (**Fig. 1A, B**). Not surprisingly we observe a positive correlation between *ARL13B* and *IFT88* (**Fig. 1C**). *SMO* also positively correlated with *IFT88* expression (**Fig. 1D**). However, *SHH* expression did not correlate with *ARL13B* **(FIG. 1E)** and significantly negatively correlated with *SMO, GLI2* and *IFT88* expression **(Fig. 1F-H)**.

**Figure 1.**
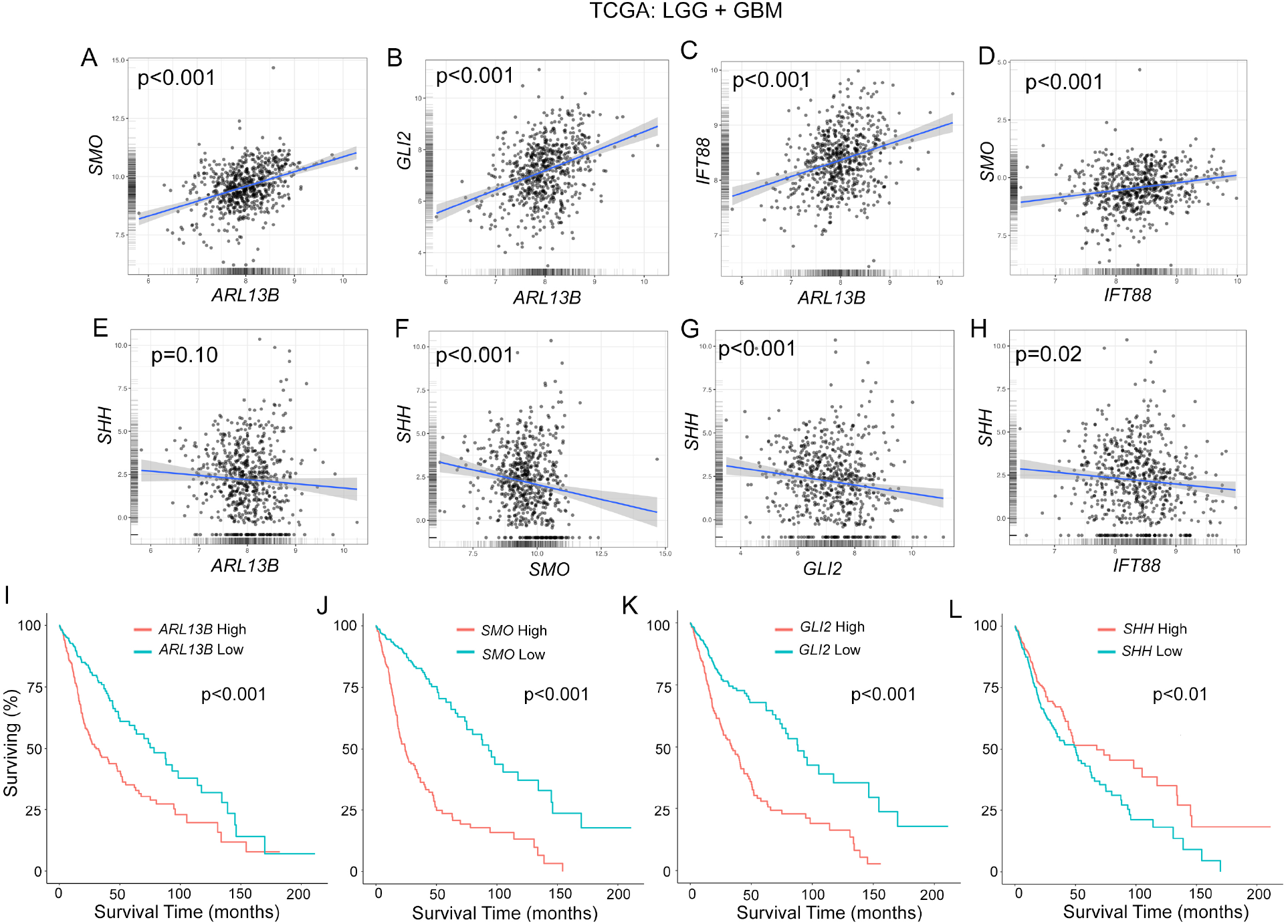
Increasing ARL13B expression correlates with SMO and GLI2 and more aggressive glioma. (**A-H**) Correlation of gene expressions for indicated gene. P values from 2-sided Pearson’s product moment correlation test are indicated. (**I-L**) Overall survival curve for the indicated gene when high (red line) or low (blue line) expressed (cutoff = median). P values represent log-rank test. All data were extracted from Gliovis database using TCGA for low and high grade glioma.

These correlations are also reflected in overall survival curves of glioma patients. Higher expression of *ARL13B* (**Fig. 1I**), *SMO* (**Fig. 1J**), *GLI2* (**Fig. 1K**) correlate with significantly shorter survival outcomes. In contrast, higher *SHH* expression correlates with significantly longer survival (**Fig. 1L**). These data suggest increased *ARL13B* is accompanied by *SMO* and *GLI2*, but not *SHH* which is surprising as it may be predicted to accompany changes associated with the SHH pathway.

### Increasing functional ARL13B in glioma cilia elongates them and promotes SMO accumulation

We next examined the effects of increasing ARL13B on glioma cilia morphology and SMO localization. To test whether ARL13B stimulation of ciliary lengthening and SMO accumulation depends on ARL13B levels and function within the cilium, we took advantage of known mutations in ARL13B. The T35N mutation ablates the GEF function and alters the GxP binding of ARL13B but localizes to cilia (Ivanova et al., 2017; Lu et al., 2015), whereas V358A mutation has normal GEF activity, but is undetectable in cilia (Gigante et al., 2020; Mariani et al., 2016). Previous studies showed that increasing transfected amounts of *ARL13b:GFP* cDNA progressively increases protein and cilia length in zebrafish embryos (Lu et al., 2015), which we confirmed in our cultures (**Suppl. Fig 2a**). Thus, we transfected parental S7 cells with increasing concentrations of GFP-tagged WT, T35N, and V358A plasmid cDNA and immunostained cells for GFP, acetylated alpha tubulin (aaTUB), or ARL13B. On GFP+ transfected cells, we measured the length of GFP^+^ cilia (for WT and T35N) or aaTub^+^/ARL13B^+^ cilia (for V358A). We found that WT ARL13B:GFP, but not T35N or V358A mutants, significantly elongated cilia as cDNA concentrations increased (**Fig. 2A**). We also confirmed that after immunostaining for ARL13B, cells transfected with WT and T35N displayed significantly increased total ciliary ARL13B compared to untransfected cells/cilia (**Suppl Fig. 2B**). We then generated an S7 cell line expressing ARL13B^WT^:GFP and ARL13B^V358A^:GFP using piggybac transgenesis. As expected, GFP expression in ARL13B ^WT^:GFP cells was localized to plasma membranes and cilia (**Fig. 2B, D(d1)**, **Suppl. Fig. 2C**), whereas in ARL13B^V358A^:GFP cells, GFP was largely undetectable in cilia but expressed throughout the plasma membranes (**Fig. 2C, D (d2), Suppl Fig. 2C**). Strikingly, immunostaining revealed significant accumulation of SMO in ARL13B^WT^:GFP cilia tips (**Fig. 2B, D(d1)**), consistent with previous studies in L0 cells (Hoang-Minh et al., 2018), that did not occur in ARL13B^V358A^:GFP cilia tips (**Fig. 2C, D(d2)**). The *ARL13B*^*V358A*^*:GFP* mutant suggests ARL13B inside cilia is stimulating SMO accumulation. Thus we examined SMO expression while increasing expression of the ARL13B^T35N^:GFP mutant, however SMO did not accumulate in these mutant cilia (**Suppl. Fig. 2D-F**).

**Figure 2.**
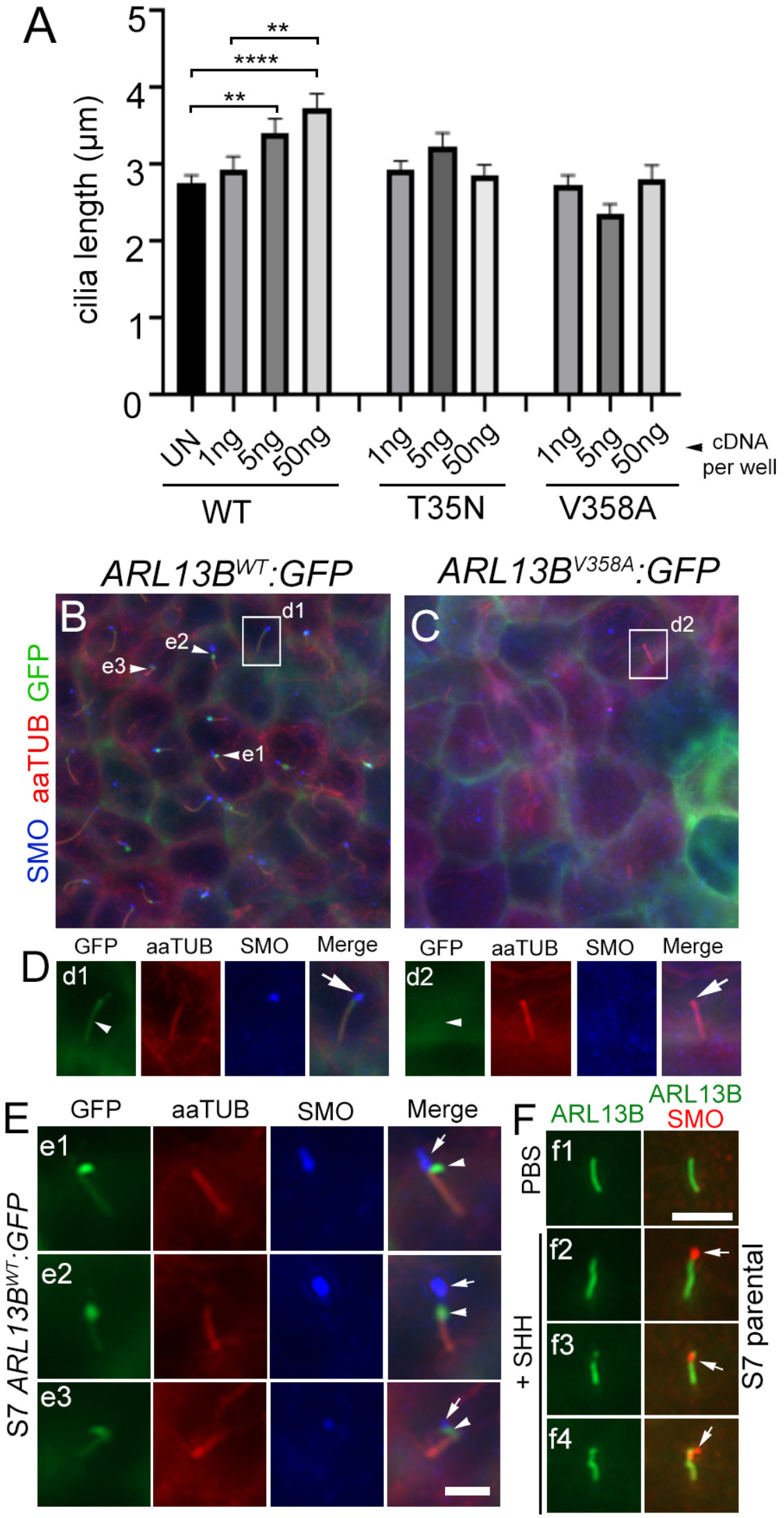
Increasing ciliary ARL13B promotes ciliary SMO enrichment glioma cilia. (**A**) Parental S7 cells were transfected with indicated concentration of ARL13B^WT^:GFP, ARL13B^T35N^:GFP, or ARL13B^V358A^:GFP cDNA. Lengths of GFP^+^ cilia (n= 22-47 per group) were measured after 48hr and compared to untransfected (UN) endogenous ARL13B^+^ cilia (n=34). **p<0.01, ****p<0.001 (ANOVA). (**B**,**C**) S7 transgenic cell lines expressing ARL13B^WT^:GFP (**B**) or ARL13B^V358A^:GFP (**C**). GFP+ cells were immunostained for acetylated alpha-tubulin (aaTUB, red) and Smoothened (SMO; blue). (**D**) Zoom of boxed areas in B,C. ARL13B^WT^:GFP^+^ cilium (arrowhead, **d1**) colocalizes with aaTUB and SMO puncta (arrow). The ARL13B^V358A^:GFP^+^ cilium is positive for aaTUB but lacks GFP (arrowhead) and SMO puncta (arrow). (**E**) Three examples of ciliary tips in ARL13B^WT^:GFP in which the GFP and SMO signals dissociate. (**F**) Cilia of parental S7 cells treated with vehicle (PBS) (**f1**) or recombinant human SHH [0.1ug/μl] (**f2-f4**) and fixed after 24hr. SHH-treated cilia display SMO puncta (arrow) that lack or weakly expressing ARL13B (arrowhead). Scale bars in E, F = 5μm.

Closer examination of ARL13B^WT^:GFP expressing cell cilia suggests SMO dissociates from ARL13B at the distal cilia tips giving the appearance that it may be budding off of the cilium (**Fig. 2E**). This phenotype is unlikely an overexpression artifact because similar patterns of distal tip puncta that are strongly SMO+, but weakly labeled for ARL13B, also occur in parental S7 cells stimulated with exogenous SHH (**Fig. 2F**).

Collectively, these data suggest that increasing functional ARL13B inside the cilium leads to lengthening of glioma cilia and distal tip accumulation of SMO that is independent of exogenous SHH exposure.

### Increasing ARL13B expression promotes abnormal tip accumulation of WT, activated SMO

We next asked if increasing ARL13B promotes and accelerates a flag-tagged form of SMO to glioma ciliary distal tips. First, we transfected two different parental glioma cell lines (L0 and S7) with FLAG-SMO which revealed SMO localized along the length of endogenous ARL13B (**Fig. 3A, B**). However, in the context of ARL13B^WT^:GFP overexpression, SMO-FLAG accumulated at the distal tips (**Fig. 3C**), similar to the endogenous SMO staining in transgenic lines (Fig. 2b,d,e). To determine if increasing ARL13B enhances distal tip accumulation of SMO, we transfected low (5ng/well) or high (200ng/well) concentrations of *ARL13b*^*WT*^*:GFP* plasmid while maintaining a low concentration of FLAG-Smo plasmid (10ng/well) and stained cells for GFP, FLAG and PCM1 to label the pericentriolar material around the ciliary basal body. Cells transfected with 5ng/well of ARL13B:GFP showed FLAG/SMO label that coincided with ARL13B^WT^:GFP (**Fig. 3D**). However, at 200ng/well the FLAG/SMO label began accumulating at the distal tips of ARL13B*WT*:GFP^+^ cilia (**Fig. 2E, compare arrowheads to arrows**). These results suggest increasing functional ARL13B in cilia drives SMO toward the ciliary tip.

**Figure 3.**
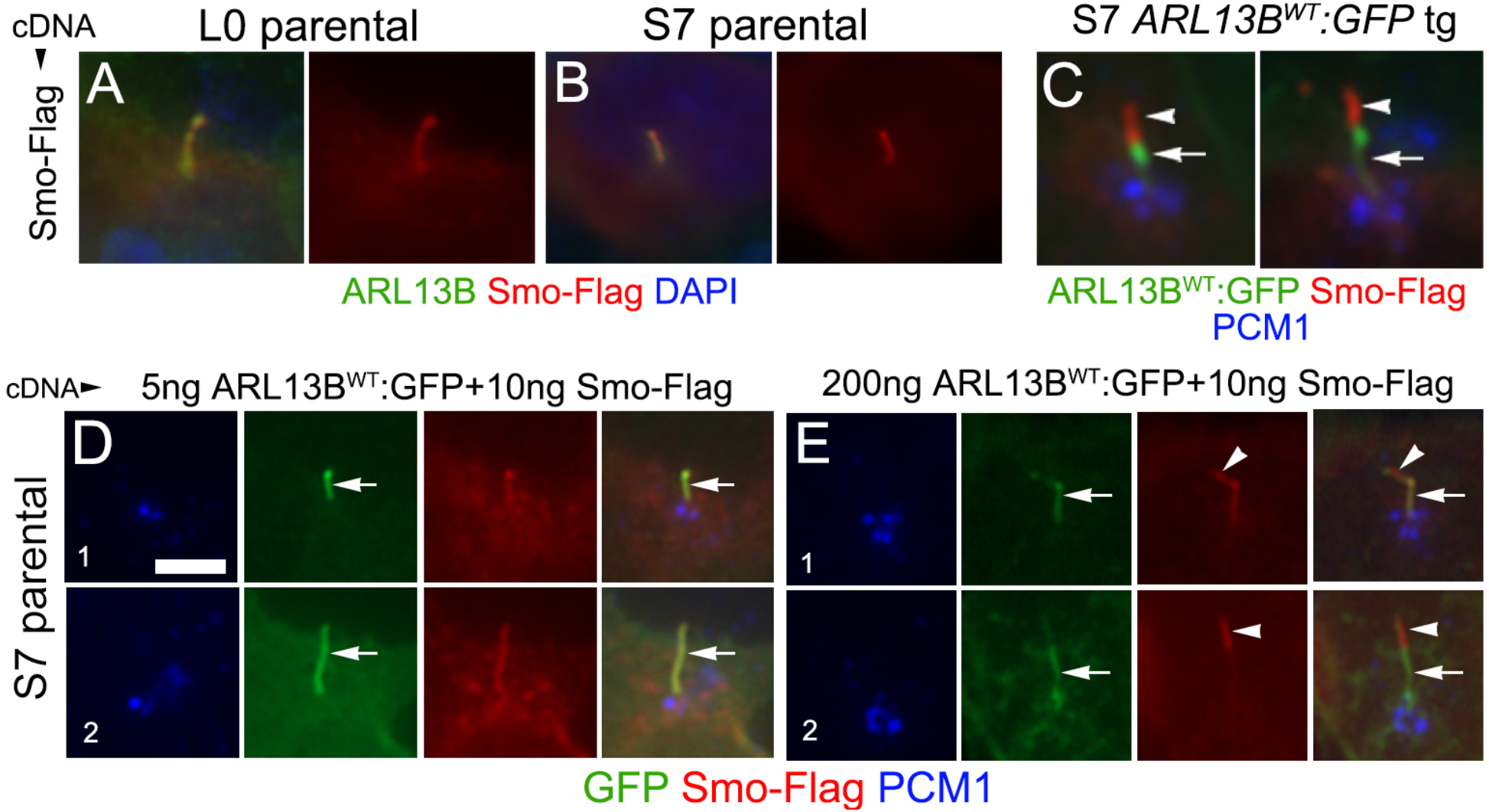
Increasing ARL13B expression stimulates ciliary tip accumulation of Flag-tagged Smo. **A, B**) L0 and S7 parental cilium expressing wildtype flag-tagged Smo (red) which co-localizes along the length of endogenous ARL13B^+^ cilia (arrows). **C**) S7 ARL13b:GFP^+^ cilia transfected with flag-Smo shows abnormal Smo accumulation (arrowheads) at the tips of cilia. Immunolabeling for PCM1 reveals the pericentriolar material that usually concentrates around the ciliary base (blue). **D**,**E**) Increasing ARL13B:GFP cDNA while holding the amount of Smo-Flag constant shows Smo/Flag accumulation at the tip of a cilium transfected with higher ARL13B:GFP cDNA. cDNA amounts are ng/well. Scale bar (D)=5μm.

It is unclear if the accumulated SMO in glioma cilia in ARL13B overexpressing cells is in an active or inactive state. To address the role of ARL13B in promoting active SMO, we used Hh-responsive mouse NIH3T3 cells (Kim et al., 2009). Previous studies showed that *GLI2* is a key mediator of activated SMO in the cilium to transcriptional changes in the nucleus (Kim et al., 2009; Santos and Reiter, 2014). We treated NIH3T3 cells with recombinant human SHH or the SMO agonist, SAG, and found SMO and GLI2 enriched in cilia (**Fig 4A-C,E,F**). Cyclopamine inhibits SMO activation yet promotes SMO ciliary enrichment, providing a way to uncouple SMO localization and activation (Kim et al., 2009; Wang et al., 2009). Indeed, we confirmed that cyclopamine stimulated SMO into cilia (**Fig 4D,G**), but we did not detect GLI2 (**Fig. 4D,H**). Taken together, these data indicate that coordination of SMO and GLI2 ciliary enrichment is due to activated SMO in cilia recruiting GLI2.

**Figure 4.**
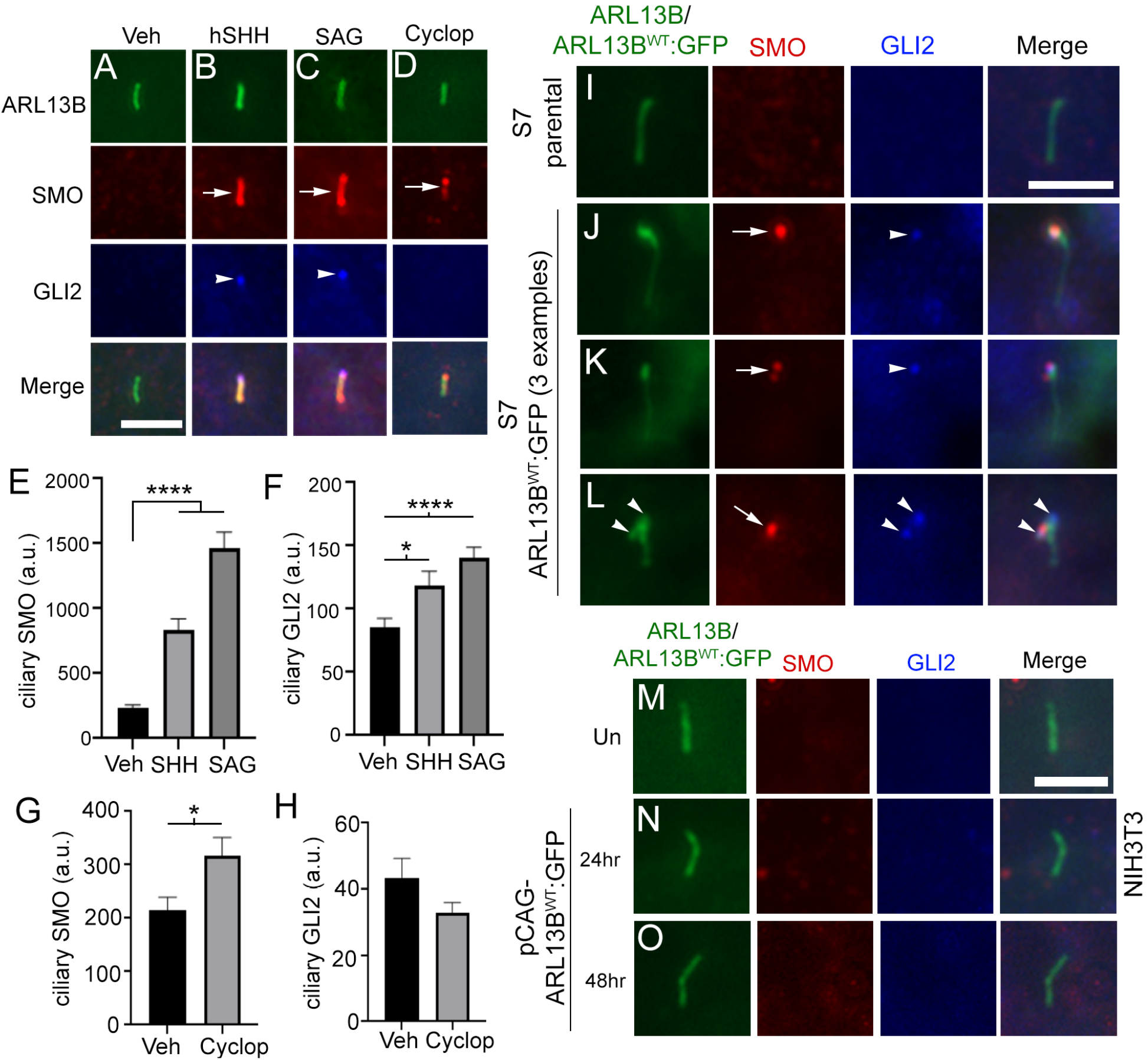
ARL13B overexpression drives active SMO and GLI2 localization in glioma but not fibroblast cilia. (**A-D**) NIH3T3 cells were treated with vehicle (Veh), 0.1μg/ml recombinant human SHH, 200nM SAG or 5μM cyclopamine (Cyclop). After 24hr, cells were fixed and triple immunolabeled for ARL13B (green), SMO (red) and GLI2 (blue). (**E**) Quantification of the SMO fluorescence intensity within cilia of untransfected (veh) (n= 39 cilia), SHH (n= 32 cilia), SAG (n=44 cilia), (**F**) Quantification of the GLI2 fluorescence intensity within cilia of untransfected (veh) (n= 39 cilia), SHH (n= 32 cilia), SAG (n=44 cilia),(**G**) Quantification of the SMO fluorescence intensity within cilia of Veh (n= 37 cilia) or cyclop (n= 42 cilia), (**H**) Quantification of the GLI2 fluorescence intensity within cilia of Veh (n= 37 cilia) or cyclop (n= 42 cilia), (**I**) S7 parental (**I**) cells immunolabeled for ARL13B (green), SMO (red) and GLI2 (blue). **J-L**) Three examples of ARL13BWT:GFP+ cilia immunolabeled for SMO and GLI2. Puncta of SMO (arrows) and GLI2 (arrowhead) are localized at the ciliary tips. (**M**) Untransfected NIH3T3 cells immunolabeled for SMO and GLI2. (N,O) NIH3T3 cells transfected with ARL13BWT:GFP cDNA for 24hr (**N**) or 48hr (**O**) and immunolabeled for SMO (red) and GLI2). Scale bars in A, I, M = 5μm. *p<0.05 ****p<0.001 (ANOVA).

Considering GLI2 localization correlated with active ciliary SMO in NIH3T3 cells, we next immunolabeled S7 parental and S7 *ARL13b*^*WT*^*:GFP* cells for SMO and GLI2.

Compared to S7 parental cilia that lack SMO and GLI2 (**Fig. 4I**), we observed ARL13B^WT^:GFP^+^ cilia whose tips were positive for both (**Fig. 4J-L**), further implicating accumulation of activated SMO. To assess whether this phenomenon is specific to glioma cells, we asked whether *ARL13b*^*WT*^*:GFP* overexpression in NIH3T3 cells also stimulates both SMO and GLI2 into cilia. However, at 24 or 48hours after transfection we did not observe increased SMO or GLI2 in ARL13bB:GFP+ cilia in this cell line (**Fig. 4M-O**). Altogether these data support the likelihood that increased ARL13B promotes ciliary enrichment of activated SMO and GLI2, and that there may be cell type specificity to the effects of increasing ARL13B on SMO/GLI2 trafficking.

### ARL13B-induced accumulation of ciliary SMO and GLI2 is resistant to potent SMO inhibitors

If ARL13B can drive SMO/GLI into cilia, we asked if this can be disrupted with current SMO inhibitors. As we show above and reported by others, cyclopamine promotes ciliary localization of SMO, potentially because the drug alters the conformation of SMO such that it cannot be ubiquitinated and removed from cilia(Desai et al., 2020; Kim et al., 2009; Wang et al., 2009). Therefore, to test whether increased ciliary SMO is due to ARL13B overexpression, we sought out other SMO inhibitors that disrupt SMO ciliary localization. In NIH3T3 cells, GDC-0449 (Vismodegib) has been shown to block the SHH-induced increase in ciliary SMO (Peluso et al., 2014). We confirmed that SHH-stimulated ciliary SMO in S7 parental glioma cells was blocked by pretreatment with GDC-0449 (**Fig. 5A-H**). Despite exposure to different concentrations of GDC-0449, we found that ARL13B^WT^:GFP^+^ cilia remain able to accumulate ciliary SMO at their distal tip in the absence of exogenous SHH (**Fig. 5I-L**). Treatment with 5μM GDC-0449 also does not block accumulation of GLI2 with SMO at the distal tips of ARL13B^WT^:GFP^+^ cilia (**Fig. 5 M,N**). Similarly, SMO and GLI2 accumulation was observed after 24hr treatment with 5μM cyclopamine (**Fig. 5O**). These results suggest ARL13B-driven changes in ciliary SMO are resistant to potent pharmacological inhibitors of SMO in glioma cells.

**Figure 5.**
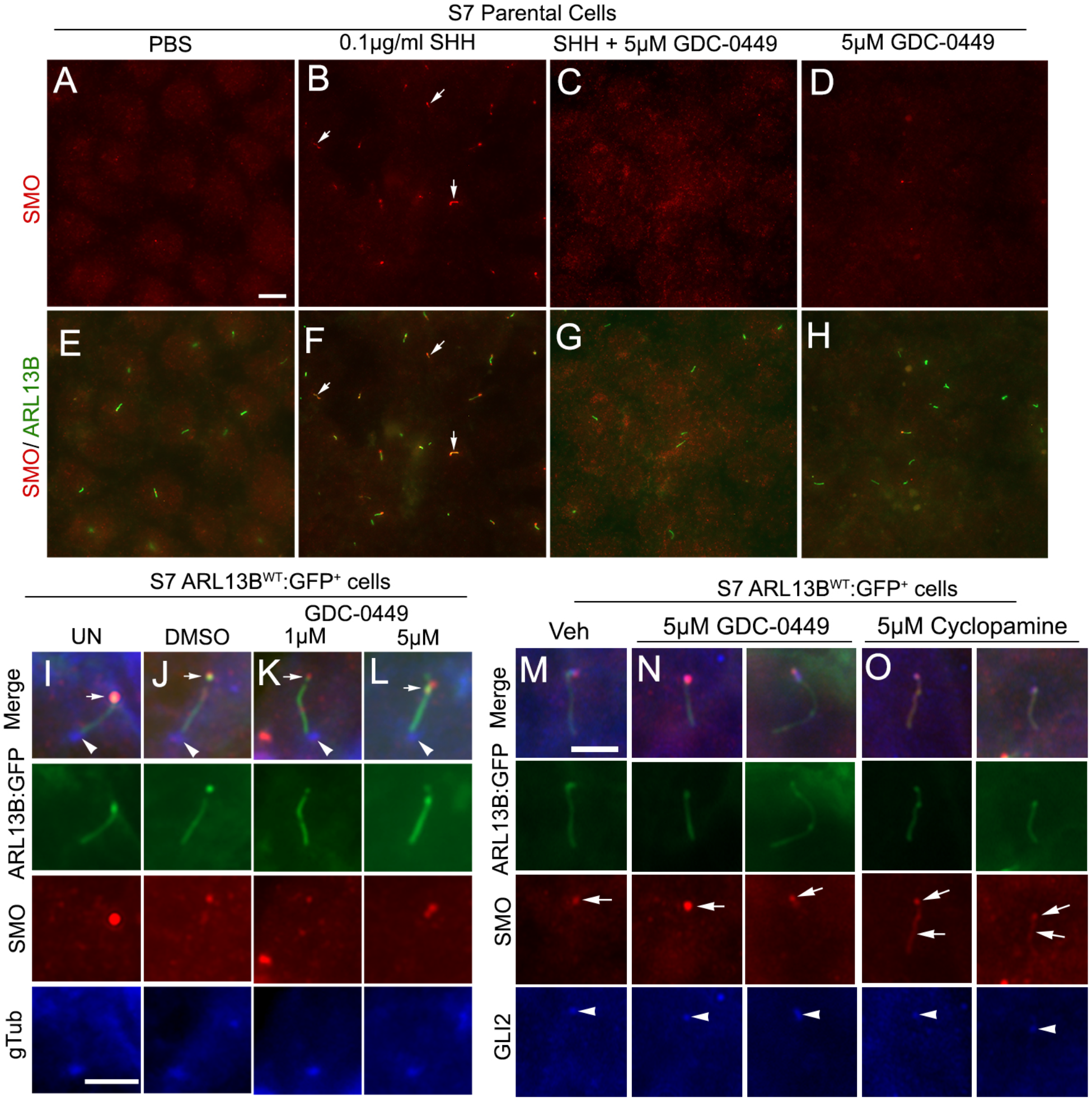
GDC-0449 (Vismodegib) and cyclopamine do not block the ARL13B:GFP-induced increase in ciliary SMO and GLI2. (**A-H**) S7 parental cells treated with PBS (**A**,**E**), 0.1ug/ml rhSHH (**B, F**), rhSHH+5μM GDC-0449 (**C**,**G**), or 5μM GDC-0449 (**D**,**H**). After 24hr, cells were fixed and immunolabeled for SMO (red) and ARL13B (green). SHH treatment induces a robust increase in ciliary SMO (arrows in **B**,**F**) in ARL13B+ cilia but not in GDC-0449 treated cells. (**I-L**) S7 ARL13B^WT^:GFP^+^ cells treated with 1 or 5μM GDC-0449 display SMO (red) at their cilia tips (arrows). The cilia base is immunolabeled with gamma-tubulin (gTub, blue, arrowheads). **M-O**) S7 ARL13B^WT^:GFP^+^ cells treated with veh (**M**), 5μM GDC-0449 (**N**, 2 examples), or 5μM cyclopamine (**O**, 2 examples), fixed after 48hr and immunolabeled for SMO (red) and GLI2 (blue). The cilia/ciliary tips were SMO+ (arrow) and GLI2+ (arrowheads) in all groups. Scale bars (in μm) in A =10, I =5 M =5.

### TMZ chemotherapy stimulates SMO and GLI2 into ARL13B^+^ cilia

The chemotherapeutic TMZ is reported to increase the transcriptional and translational expression of ARL13B (27). It is also accompanied by an increase the number of ARL13B^+^ cilia, and a potential mechanism that promotes resistance to the therapy (Shi et al., 2022; Shireman et al., 2021; Wei et al., 2022). We asked if the TMZ-induced ARL13B changes are accompanied by changes in ciliary SMO or GLI2 in our glioma cells. In vitro studies often require high concentrations (100μM to 4000μM) of TMZ to begin eliciting cell death (Herbener et al., 2020). We treated parental S7 cells to vehicle or 100μM TMZ for 24 hours, fixed and immunostained cells for endogenous ARL13B, SMO and GLI2. We observed enriched SMO and GLI2 in ARL13B^+^ cilia of TMZ-treated cells (**Fig. 6A,B**). Quantification of the percent of ARL13B^+^ in our cultures (**Fig. 6C**) and ciliary intensity of SMO (**Fig. 6D**) and GLI2 (**Fig. 6E**) revealed significant increases after TMZ treatment. To assess the cell type specificity of this effect, we examined the same parameters using NIH3T3 cells. TMZ treatment led to fewer ARL13B^+^ ciliated 3T3 cells (**Fig. 6F**), no significant change in ciliary SMO intensity (**Fig. 6G**) and significantly reduced intensity of ciliary GLI2 (**Fig. 6H**). Thus, TMZ effects on SMO and GLI2 in ARL13B^+^ cilia differ between glioma and a non-cancer cell type.

**Figure 6.**
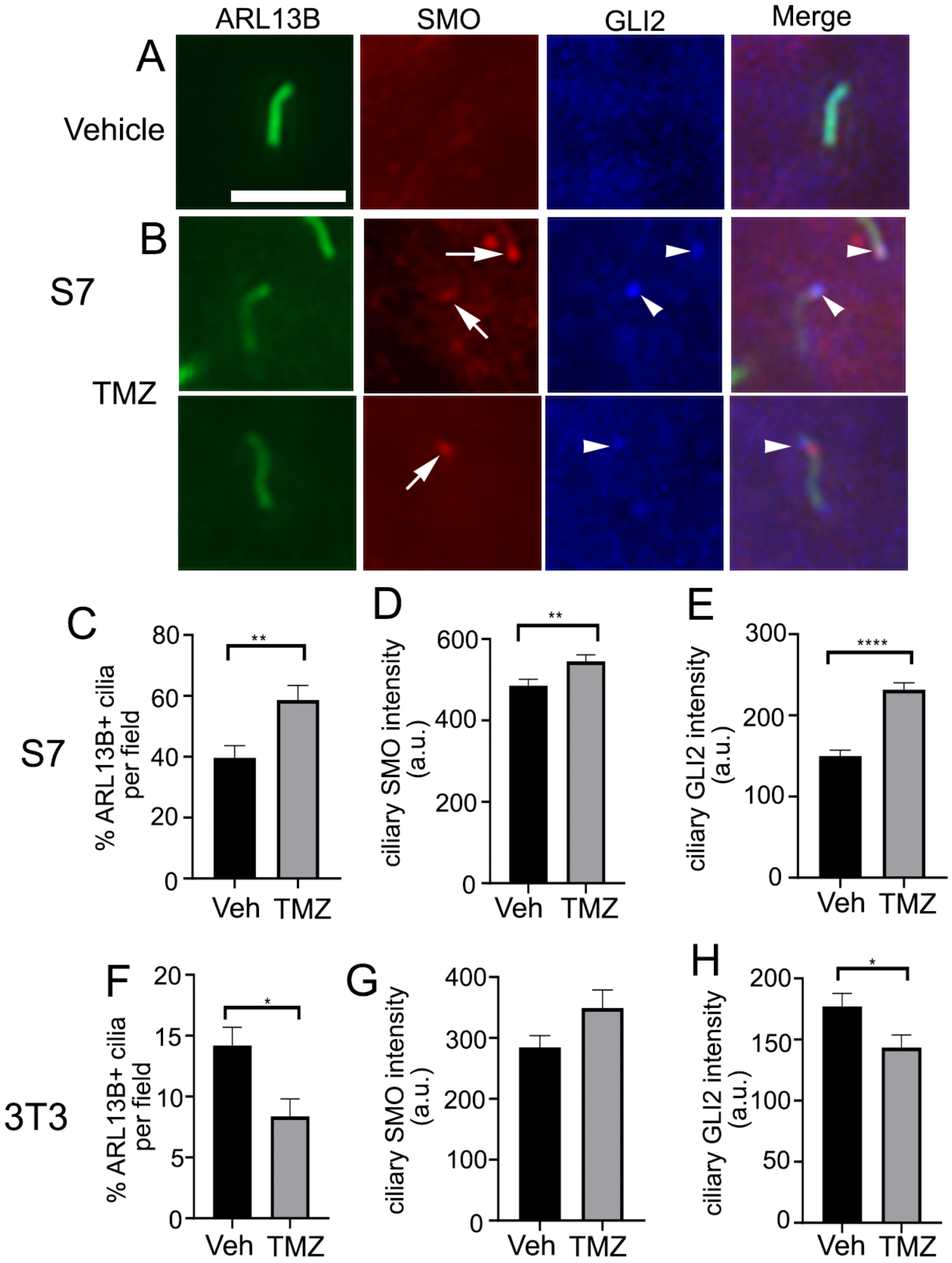
Temozolomide stimulates SMO and GLI2 into ARL13B+ cilia of S7 glioma but not NIH3T3 cells. (**A**,**B**) S7 parental cells treated with vehicle (**A**) or 100μM TMZ (**B**, 2 examples) fixed after 24hr, and immunolabeled for ARL13B, SMO (red) and GLI2 (blue). In TMZ treated cells, arrows point to SMO+ cilia with GLI2+ cilia tips (arrowheads). (**C-E**) Bar graphs show the percent of ciliated cells (**C**), ciliary SMO intensity (a.u. = arbitrary units) (**D**), or ciliary GLI2 intensity (**E**) for S7 parental cells after Veh or TMZ treatment. (**F-H**) Bar graphs show the percent of ciliated cells (**F**), ciliary SMO intensity (a.u. = arbitrary units) (**G**), or ciliary GLI2 intensity (**H**) for NIH3T3 cells after Veh or TMZ treatment. *p<0.05, **p<0.01, ****p<0.001. Bar = 5μm.

## Discussion

We find that increasing ARL13B in glioma cilia stimulates ciliary elongation and an increase in SMO/GLI2 accumulation. This appears to be due to a function of ARL13B within cilia as we see no changes in cilia or SMO/GLI2 accumulation when we increased cellular ARL13B. The ciliary ARL13B-driven increase in SMO is likely an active form of SMO that recruits GLI2, resembling cells that have been stimulated exogenously with SHH or SAG. The ARL13B-driven increase in SMO and GLI2 appears resistant to SMO inhibitors, and is observed after TMZ chemotherapy. Our findings raise the possibility that increasing ARL13B beyond a certain level inside the cilium alters the biology within glioma cilia leading to SMO/GLI accumulation (**Fig. 7**). Considering our previous studies which showed glioma cells excise vesicles from their distal tips(Hoang-Minh et al., 2018), it is possible the accumulation of active SMO and GLI exit from cilia in vesicles may be capable of intercellular signaling to other cancer or unknown recipient host cells, though this latter phenomenon requires further exploration.

**Figure 7.**
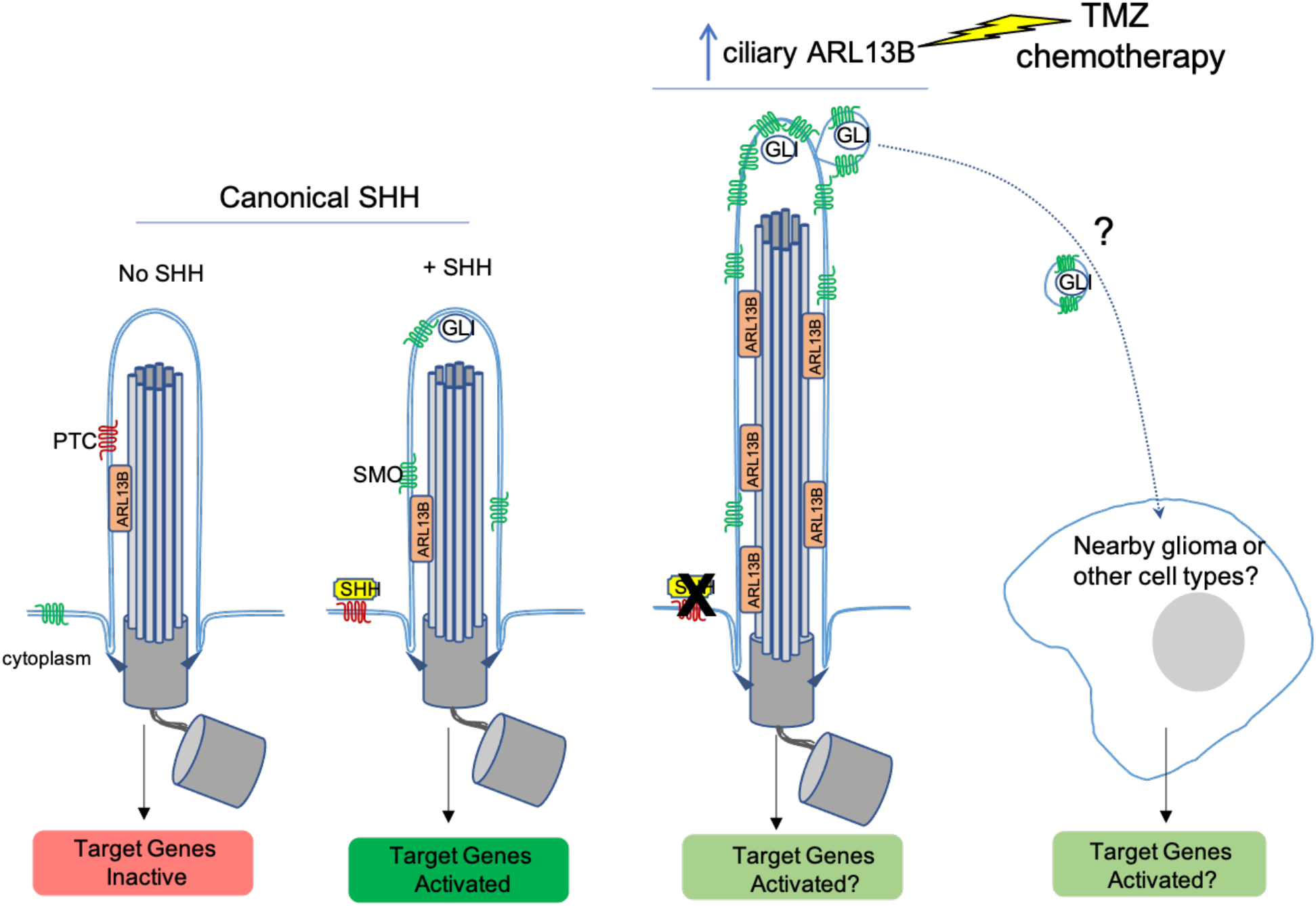
Model comparing SHH vs increasing ARL13B on the localization of SMO/GLI. Left: Canonical SHH pathway activation involving SHH induced change in ciliary SMO. Right: in absence of SHH, increasing ARL13B in cilia elongates the structure and alters cilia membrane protein components.

### ARL13B-driven increases in ciliary SMO: failed retrieval of activated SMO or a resistance mechanism in glioma?

In cells with increased ARL13B expression there was significant ciliary distal tip accumulation of SMO (Figs. 2 and 3). One possibility is that ARL13B somehow drives activated SMO into cilia faster than it can be brought back out of cilia. In IMCD3 cells, activated GPCRs that are not retrieved by intraflagellar transport and BBSome trains are ectocytosed as ciliary vesicles (Nager et al., 2017; Ye et al., 2018). Thus, ARL13B increases in the cilia may enhance SMO accumulation by disrupting the ability of IFT machinery to retrieve activated SMO. The initial front loading of SMO into cilia may be related to reports in gastric tumor cells that ARL13B directly binds and stabilizes SMO (Shao et al., 2017). However, we do not know if this potential complex during ciliary entry exists in glioma. If it does exist, it appears to dissociate at the distal tip based on our immunolabeling of distal tip buds that appear strongly SMO but weakly ARL13B-labeled (Fig. 2), which would suggest other factors are present in the glioma ciliary tip to promote this dissociation. Alternatively, in glioma ARL13B does not bind or promote SMO ciliary localization. In addition, it was surprising that we did not find the same ARL13B-induced SMO accumulation in cilia of NIH3T3 cells, which have constitutive SHH/SMO signaling. Thus, glioma cells may have other changes that are more permissive to SMO trafficking.

The ARL13B driven increases in ciliary SMO and downstream pathway activation appear resistant to both cyclopamine (Shao et al., 2017) and GDC-0449 (Fig. 5), and increasing ARL13B somehow overrides the efficacy of these inhibitors. It is not clear if ARL13B changes properties of the ciliary membrane such that SMO inhibitors lose their efficacy and stimulate SMO entry. For example, SMO is activated in cilia by membrane phospholipids (oxysterols) uniquely concentrated in ciliary plasma membrane (Raleigh et al., 2018). Increases in ARL13B could alter the organization of these phospholipids since ARL13B is reported to recruit the lipid phosphatase, inositol polyphosphate-5-phosphatase E (INPP5e) (Humbert et al., 2012). Alternatively, SMO entry into cilia is also promoted by cAMP-PKA pathway stimulation (Milenkovic et al., 2009). However, SMO is believed to be Gi coupled receptor that decreases cAMP (Ogden et al., 2008; Riobo et al., 2006; Shen et al., 2013). In the context of ARL13B overexpression, it could promote entry of other ciliary receptors that stimulate cAMP-PKA and indirectly, additional SMO. Whatever the mechanism, ARL13B could change the ciliary membrane or other GPCRs in the cilium that favor SMO entry/activation and reduces inhibitor efficacy.

The ARL13B-linked changes in SMO and GLI2 can also be observed without direct overexpression. Recent studies showed that TMZ stimulates an increase in ARL13B expression, including the length and number of cilia (Shi et al., 2022; Shireman et al., 2021; Wei et al., 2022). Consistent with these results, we found TMZ stimulated ARL13B+ ciliogenesis and SMO/GLI2 enrichment (Fig. 7), and that this effect was not observed in NIH3T3 cells. The treatment-related nature of these findings may resemble increases in SMO/GLI2 observed in other drug-resistant, non-small cell lung carcinoma cell lines (Jenks et al., 2018). Why SMO/GLI2 is stimulated into cilia following TMZ is unclear, but may be involved the purine synthesis pathway reported to promote chemoresistance (Shireman et al., 2021). Whether TMZ leads to release of SMO/GLI2 in the form of vesicles, which we observed during ARL13B:GFP overexpression (Hoang-Minh et al., 2018), requires further investigation.

### What does high expression of ARL13B and SMO in TCGA signify?

High *ARL13b, SMO* and *GLI2* expression correlate with lower patient survival in the TCGA (Fig. 1)(Hoang-Minh et al., 2018; Shireman et al., 2021). However, this database does not inform about the ciliation status within tumors. Our results suggest it may be worth determining if shorter patient survival correlates with higher frequencies of ARL13B+ and/or SMO+/GLI2+ cilia. In addition, high *ARL13b, SMO*, and *GLI2* expression does not align with SHH expression (Fig. 1). We show that high relative to low expression of SHH predicts longer patient survival which is counterintuitive given ARL13B, SMO and GLI2 mediate SHH signaling (Gigante et al., 2020; Larkins et al., 2011). The relationship between ARL13B, SMO and GLI2 may be independent of SHH expression in glioma. Indeed, in our previous study (Hoang-Minh et al., 2018), and in this study (Fig. 2), we find that ARL13B-mediated enrichment of ciliary SMO resembles the effects of stimulating parental cells with SHH. Alternatively, the correlation between ARL13B, SMO and GLI2 is related to other factors that possibly control *ARL13b* expression in the tumors. We and others (Shireman et al., 2021) find TMZ stimulates ARL13B. Given this observation, a significant percentage of tumors may increase ARL13B, SMO and GLI2 in response to TMZ and therefore promote more aggressive tumors. Considering ARL13B-driven SMO appears resistant to SMO inhibitors in culture it may not be surprising to observe SMO inhibitors fail to control tumorigenesis. Future studies need to identify SMO inhibitors that can block ARL13B-mediated changes in SMO, and examine whether or not TMZ drives SMO/GLI2 into cilia in in vivo models of patient tumors.

## Materials and Methods

### Cell culture

Human U138 MG (Cat# HTB-16), U87 MG (Cat# HTB-14), A172 (Cat# CRL-1620) and rat F98 (Cat# CRL-2397) cells were all obtained from ATCC and maintained in recommended media conditions. Mouse KR158 glioma cells were derived from a murine grade III anaplastic astrocytoma (Gursel et al., 2011), and maintained in DMEM ([-] sodium pyruvate; Corning; Cat #10017CV), 10% heat-inactivated fetal bovine serum (FBS) (Atlanta Biologicals; Cat #SH30070.03) and 1% penicillin–streptomycin. L0 (grade IV glioblastoma from a 43 year old male) and S7(grade II glioma from a 54 year old female with EGFR amplification) cell lines were isolated and maintained as previously described (Deleyrolle et al., 2011; Hothi et al., 2012; Lin et al., 2015; Sarkisian et al., 2014). L0 and S7 cells were grown as floating spheres and maintained in NeuroCult NS-A Proliferation medium and 10% proliferation supplement (STEMCELL Technologies; Cat# 05750 and #05753), 1% penicillin–streptomycin (Thermofisher, Cat# 15140122), 20 ng/ml human epidermal growth factor (hEGF) (Cat #78006), and 10 ng/ml basic fibroblast growth factor (bFGF) (STEMCELL Technologies; Cat #78003).

For S7 cells, the media was supplemented with 2 μg/ml heparin (STEMCELL TechnologiesCat #07980). When cells reached confluency, or spheres reached approximately 150 μm in diameter, they were enzymatically dissociated by digestion with Accumax (Innovative Cell Technologies; Cat#AM-105) for 10 min at 37 °C. For human cells grown on glass coverslips, NeuroCult NS-A Proliferation medium was supplemented with 10% heat inactivated fetal bovine serum (FBS) (Cytiva, Cat #SH30070.03HI). NIH3T3 cells (ATCC Cat# CRL-1658) were maintained in DMEM (ATCC Cat# 30-2002), 10% FBS and 1% penicillin–streptomycin. All cells were grown in a humidified incubator at 37 °C with 5% CO_2_. Where indicated, indicated cell lines were treated with various drugs including cyclopamine (Calbiochem; Cat#239803) (diluted in dimethyl sulfoxide (DMSO)), temozolomide (TMZ) (Sigma; Cat# T2577) (diluted in DMSO), GDC-0449 (Cayman Chemical; Cat #13613), SAG (Calbiochem; Cat #566661), recombinant human SHH (high activity (R&D Systems: Cat # 8908-SH-005/CF), and fixed after indicated treatment durations and processed as described below. Where indicated, cells were also transiently transfected with pFlag-Smo (PD22) (gift from G. Pazour), pCAG-Ar13b^WT^:GFPpb, pDest-Arl13b^T35N^:GFP cDNA using Lipofectamine 3000(Invitrogen; cat # L3000-008), and fixed at the indicated timepoints as described below.

S7 clones stably expressing mouse *Arl13B*^*WT*^*:GFP* or *Arl13B*^*V358A*^*:GFP* were generated using piggybac transposase similar to previously described(Hoang-Minh et al., 2018). Briefly S7 cells were grown as adherent in 6-well plates and transfected at 60-70% confluence with a total of 500 ng per well of pCAG-pBase and pCAG-Arl13b (WT):GFPpb or pCAG-ARL13B(V358A):GFP vectors using 10μl of Lipofectamine 3000. To generate the pCAG-Ar13b^WT^:GFPpb or pCAG-Ar13b^V358A^:GFPpb vector, we subcloned the C-terminal GFP-tagged Arl13b sequence from pDest-Arl13b WT:GFP or pDest-Arl13b^V358A^:GFP into the pCAG-pb vector. Approximately 1 week after transfection, individual GFP^+^ clones were sorted and expanded in 96-well plates each containing 250 μL of supplemented S7 medium without FBS using a BD FACS Aria II Cell Sorter (BD Biosciences, San Jose, CA). Cell debris were excluded from the analysis by forward- and side-scatter gating. Subsequently expanded GFP^+^ clones were propagated.

### Western Blot

Western blot was performed as recently described (Shi et al., 2021). Briefly, cells were harvested at indicated time points and lysed in 1× radioimmunoprecipitation assay (RIPA) buffer (Cell Signaling, Danvers, MA, USA; Cat# 501015489) containing 1× protease inhibitor cocktail (Sigma, St. Louis, MO, USA; Cat# P2850), phosphatase inhibitor cocktails 1 (Sigma, St. Louis, MO, USA; Cat# P5726), and 2 (Sigma; Cat# P0044), and 1× phenylmethanesulfonyl fluoride (Sigma, St. Louis, MO, USA; Cat# 93482). A total of 25-30 μg of total protein lysate per lane were separated on a 4–12% Bis-Tris gel (Thermofisher; Cat# NP0050). Proteins were blotted onto PVDF membranes using iBlot (program 3 for 8 min; Invitrogen, Carlsbad, CA, USA). Blots were blocked in 5% nonfat dry milk (NFDM) or bovine serum albumin (BSA, Jackson Immuno Research, West Grove, PA, USA; Cat# NC9871802) in 1× tris-buffered saline (TBS) with 0.1% Tween (TBST) for 20 min and then incubated in primary antibodies in 2.5% NFDM or BSA in 1× TBST for 24 h at 4 °C. Blots were rinsed and probed in the appropriate horseradish peroxidase (HRP)-conjugated secondary antibody (1:10,000; BioRad, Hercules, CA, USA) for 30 min at RT in 2.5% NFDM or BSA in 1× TBST. Finally, blots were rinsed in 1× TBS and developed using an Amersham ECL chemiluminescence kit (Global Life Sciences Solutions USA, Marlborough, MA, USA), and images were captured using an AlphaInnotech Fluorchem Q Imaging System (Protein Simple, San Jose, CA, USA). Selected areas surrounding the predicted molecular weight of the protein of interest were extracted from whole blot images.

### Immunocytochemistry

For immunocytochemical analyses, samples were fixed at indicated timepoints with 4% paraformaldehyde in 0.1 M phosphate buffer (4% PFA) for 15 min and washed with 1x PBS). Cells were immunolabeled for the indicated primary antibodies (Table 1).

**Table 1.**
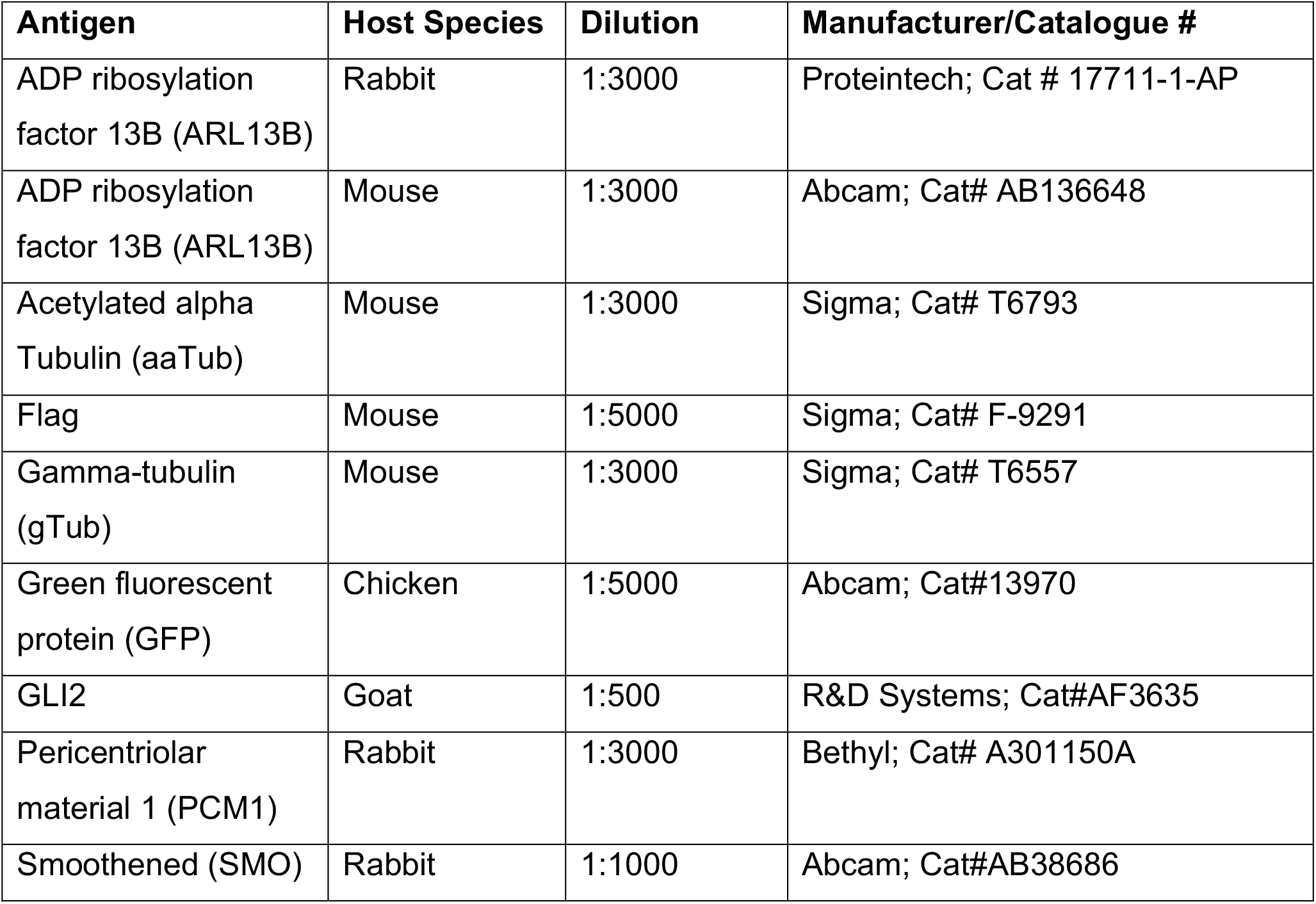
Primary antibodies used in this study.

Samples were incubated in blocking solution containing 5% normal donkey serum (NDS) (Jackson Immunoresearch; Cat#NC9624464) and 0.2% Triton-X 100 in 1x PBS for 1 hour and then incubated in primary antibodies with 2.5% NDS and 0.1% Triton-X 100 in 1x PBS either for 2 hours at room temperature (RT) or overnight at 4°C. Appropriate FITC-, Cy3- or Cy5-conjugated secondary antibodies (1:1000; Jackson ImmunoResearch) in 2.5% NDS with 1x PBS were applied for 1-2 hour at RT, and coverslips were mounted onto Superfrost™ Plus coated glass slides (Fisher Scientific, cat # 12-550-15) in Prolong Gold antifade media containing DAPI (Thermofisher; Cat# P36935). Stained coverslips were examined under epifluorescence using an inverted Zeiss AxioObserver D1 microscope using a Zeiss 40×/0.95 plan Apochromat air objective or a Zeiss 63X/1.4 plan Apochromat oil objective. Images were captured and analyzed using Zeiss ZEN software.

For analyses of cilia length, ARL13B^+^ cilia were traced using a line measurement tool in Zeiss ZEN software. For analyses of immunofluorescence staining intensity, areas were traced around the cell cilium and the mean fluorescence intensity (MFI) was background corrected. The background MFI, measured from an area between the cells, was subtracted from the cilia MFI. We analyzed cilia from at least 2-3 coverslips per group.

## Acknowledgements

We would like to thank G. Pazour (Univ. Massachusetts Medical School, Worcester MA) for Smo expression vectors. Funding for this work was supported by an American Brain Tumor Association Discovery Grant (DG1800017) supported by an Anonymous Family Foundation (to MRS).

## References

Aiman, W., and A. Rayi. 2022. Low Grade Gliomas. In StatPearls, Treasure Island (FL).

Alvarez-Satta, M., and A. Matheu. 2018. Primary cilium and glioblastoma. Ther Adv Med Oncol. 10:1758835918801169.

Bangs, F., and K.V. Anderson. 2017. Primary Cilia and Mammalian Hedgehog Signaling. Cold Spring Harb Perspect Biol. 9.

Bar, E.E., A. Chaudhry, M.H. Farah, and C.G. Eberhart. 2007a. Hedgehog signaling promotes medulloblastoma survival via Bc/II. Am J Pathol. 170:347–355.

Bar, E.E., A. Chaudhry, A. Lin, X. Fan, K. Schreck, W. Matsui, S. Piccirillo, A.L. Vescovi, F. DiMeco, A. Olivi, and C.G. Eberhart. 2007b. Cyclopamine-mediated hedgehog pathway inhibition depletes stem-like cancer cells in glioblastoma. Stem Cells. 25:2524–2533.

Caspary, T., C.E. Larkins, and K.V. Anderson. 2007. The graded response to Sonic Hedgehog depends on cilia architecture. Dev Cell. 12:767–778.

Chen, L., X. Xie, T. Wang, L. Xu, Z. Zhai, H. Wu, L. Deng, Q. Lu, Z. Chen, X. Yang, H. Lu, Y.G. Chen, and S. Luo. 2022. ARL13B promotes angiogenesis and glioma growth by activating VEGFA-VEGFR2 signaling. Neuro Oncol. noac245.

Clement, V., P. Sanchez, N. de Tribolet, I. Radovanovic, and A. Ruiz i Altaba. 2007. HEDGEHOG-GLI1 signaling regulates human glioma growth, cancer stem cell self-renewal, and tumorigenicity. Curr Biol. 17:165–172.

Deleyrolle, L.P., A. Harding, K. Cato, F.A. Siebzehnrubl, M. Rahman, H. Azari, S. Olson, B. Gabrielli, G. Osborne, A. Vescovi, and B.A. Reynolds. 2011. Evidence for label-retaining tumour-initiating cells in human glioblastoma. Brain. 134:1331–1343.

Desai, P.B., M.W. Stuck, B. Lv, and G.J. Pazour. 2020. Ubiquitin links smoothened to intraflagellar transport to regulate Hedgehog signaling. J Cell Biol. 219.

Gigante, E.D., M.R. Taylor, A.A. Ivanova, R.A. Kahn, and T. Caspary. 2020. ARL13B regulates Sonic hedgehog signaling from outside primary cilia. Elife. 9: :e50434.

Goetz, S.C., and K.V. Anderson. 2010. The primary cilium: a signalling centre during vertebrate development. Nat Rev Genet. 11:331–344.

Goetz, S.C., P.J. Ocbina, and K.V. Anderson. 2009. The primary cilium as a Hedgehog signal transduction machine. Methods Cell Biol. 94:199–222.

Goranci-Buzhala, G., A. Mariappan, L. Ricci-Vitiani, N. Josipovic, S. Pacioni, M. Gottardo, J. Ptok, H. Schaal, G. Callaini, K. Rajalingam, B. Dynlacht, K. Hadian, A. Papantonis, R. Pallini, and J. Gopalakrishnan. 2021. Cilium induction triggers differentiation of glioma stem cells. Cell Rep. 36:109656.

Gruber Filbin, M., S.K. Dabral, M.F. Pazyra-Murphy, S. Ramkissoon, A.L. Kung, E. Pak, J. Chung, M.A. Theisen, Y. Sun, Y. Franchetti, D.S. Shulman, N. Redjal, B. Tabak, R. Beroukhim, Q. Wang, J. Zhao, M. Dorsch, S. Buonamici, K.L. Ligon, J.F. Kelleher, and R.A. Segal. 2013. Coordinate activation of Shh and PI3K signaling in PTEN-deficient glioblastoma: new therapeutic opportunities. Nat Med. 19:1518–1523.

Gursel, D.B., Y.S. Connell-Albert, R.G. Tuskan, T. Anastassiadis, J.C. Walrath, J.J. Hawes, J.C. Amlin-Van Schaick, and K.M. Reilly. 2011. Control of proliferation in astrocytoma cells by the receptor tyrosine kinase/PI3K/AKT signaling axis and the use of PI-103 and TCN as potential anti-astrocytoma therapies. Neuro Oncol. 13:610–621.

Han, Y.G., and A. Alvarez-Buylla. 2010. Role of primary cilia in brain development and cancer. Curr Opin Neurobiol. 20:58–67.

Herbener, V.J., T. Burster, A. Goreth, M. Pruss, H. von Bandemer, T. Baisch, R. Fitzel, M.D. Siegelin, G. Karpel-Massler, K.M. Debatin, M.A. Westhoff, and H. Strobel. 2020. Considering the experimental use of temozolomide in glioblastoma research. Biomedicines. 8:151.

Hoang-Minh, L., L. Deleyrolle, N. Nakamura, A. Parker, R. Martuscello, B. Reynolds, and M. Sarkisian. 2016a. PCM1 depletion inhibits glioblastoma cell ciliogenesis and increases cell death and sensitivity to temozolomide. Transl Oncol. 9:392–402.

Hoang-Minh, L.B., L.P. Deleyrolle, D. Siebzehnrubl, G. Ugartemendia, H. Futch, B. Griffith, J.J. Breunig, G. De Leon, D.A. Mitchell, S. Semple-Rowland, B.A. Reynolds, and M.R. Sarkisian. 2016b. Disruption of KIF3A in patient-derived glioblastoma cells: effects on ciliogenesis, hedgehog sensitivity, and tumorigenesis. Oncotarget. 7:7029–7043.

Hoang-Minh, L.B., M. Dutra-Clarke, J.J. Breunig, and M.R. Sarkisian. 2018. Glioma cell proliferation is enhanced in the presence of tumor-derived cilia vesicles. Cilia. 7:6.

Hothi, P., T.J. Martins, L. Chen, L. Deleyrolle, J.G. Yoon, B. Reynolds, and G. Foltz. 2012. High-throughput chemical screens identify disulfiram as an inhibitor of human glioblastoma stem cells. Oncotarget. 3:1124–1136.

Humbert, M.C., K. Weihbrecht, C.C. Searby, Y. Li, R.M. Pope, V.C. Sheffield, and S. Seo. 2012. ARL13B, PDE6D, and CEP164 form a functional network for INPP5E ciliary targeting. Proc Natl Acad Sci USA. 109:19691–19696.

Ivanova, A.A., T. Caspary, N.T. Seyfried, D.M. Duong, A.B. West, Z. Liu, and R.A. Kahn. 2017. Biochemical characterization of purified mammalian ARL13B protein indicates that it is an atypical GTPase and ARL3 guanine nucleotide exchange factor (GEF). J Biol Chem. 292:11091–11108.

Jenks, A.D., S. Vyse, J.P. Wong, E. Kostaras, D. Keller, T. Burgoyne, A. Shoemark, A. Tsalikis, M. de la Roche, M. Michaelis, J. Cinatl, Jr., P.H. Huang, and B.E. Tanos. 2018. Primary cilia mediate diverse kinase inhibitor resistance mechanisms in cancer. Cell Rep. 23:3042–3055.

Jooma, R., M. Waqas, and I. Khan. 2019. Diffuse Low-Grade Glioma - Changing Concepts in Diagnosis and Management: A Review. Asian J Neurosurg. 14:356–363.

Kim, J., M. Kato, and P.A. Beachy. 2009. Gli2 trafficking links Hedgehog-dependent activation of Smoothened in the primary cilium to transcriptional activation in the nucleus. Proceedings of the National Academy of Sciences of the United States of America. 106:21666–21671.

Larkins, C.E., G.D. Aviles, M.P. East, R.A. Kahn, and T. Caspary. 2011. Arl13b regulates ciliogenesis and the dynamic localization of Shh signaling proteins. Molecular biology of the cell. 22:4694–4703.

Lee, D., R.C. Gimple, B.C. Prager, Z. Qiu, Q. Wu, V. Daggubati, J. Gopalakrishnan, M.R. Sarkisian, D.R. Raleigh, and J.N. Rich. 2022. Superenhancer-activation of KLHDC8A Drives Glioma Ciliation and Hedgehog Signaling. J Clin Invest. e163592.

Lin, B., H. Lee, J.G. Yoon, A. Madan, E. Wayner, S. Tonning, P. Hothi, B. Schroeder, I. Ulasov, G. Foltz, L. Hood, and C. Cobbs. 2015. Global analysis of H3K4me3 and H3K27me3 profiles in glioblastoma stem cells and identification of SLC17A7 as a bivalent tumor suppressor gene. Oncotarget. 6:5369–5381.

Liu, H., A.A. Kiseleva, and E.A. Golemis. 2018. Ciliary signalling in cancer. Nat Rev Cancer. 18:511–524.

Lu, H., M.T. Toh, V. Narasimhan, S.K. Thamilselvam, S.P. Choksi, and S. Roy. 2015. A function for the Joubert syndrome protein Arl13b in ciliary membrane extension and ciliary length regulation. Dev Biol. 397:225–236.

Mariani, L.E., M.F. Bijlsma, A.I. Ivanova, S.K. Suciu, R.A. Kahn, and T. Caspary. 2016. Arl13b regulates Shh signaling from both inside and outside the cilium. Mol Biol Cell. 27:3780–3790.

Milenkovic, L., M.P. Scott, and R. Rohatgi. 2009. Lateral transport of Smoothened from the plasma membrane to the membrane of the cilium. J Cell Biol. 187:365–374.

Moser, J.J., M.J. Fritzler, and J.B. Rattner. 2009. Primary ciliogenesis defects are associated with human astrocytoma/glioblastoma cells. BMC Cancer. 9:448.

Nager, A.R., J.S. Goldstein, V. Herranz-Perez, D. Portran, F. Ye, J.M. Garcia-Verdugo, and M.V. Nachury. 2017. An actin network dispatches ciliary GPCRs into extracellular vesicles to modulate signaling. Cell. 168:252–263 e214.

Ogden, S.K., D.L. Fei, N.S. Schilling, Y.F. Ahmed, J. Hwa, and D.J. Robbins. 2008. G protein Galphai functions immediately downstream of Smoothened in Hedgehog signalling. Nature. 456:967–970.

Omuro, A., and L.M. DeAngelis. 2013. Glioblastoma and other malignant gliomas: a clinical review. JAMA. 310:1842–1850.

Osborn, A.G., D.N. Louis, T.Y. Poussaint, L.L. Linscott, and K.L. Salzman. 2022. The 2021 World Health Organization Classification of Tumors of the Central Nervous System: What Neuroradiologists Need to Know. AJNR Am J Neuroradiol. 43:928–937.

Peluso, M.O., V.T. Campbell, J.A. Harari, T.T. Tibbitts, J.L. Proctor, N. Whitebread, J.M. Conley, K.F. White, J.L. Kutok, M.A. Read, K. McGovern, and K.L. Faia. 2014. Impact of the Smoothened inhibitor, IPI-926, on smoothened ciliary localization and Hedgehog pathway activity. PLoS One. 9:e90534.

Raleigh, D.R., N. Sever, P.K. Choksi, M.A. Sigg, K.M. Hines, B.M. Thompson, D. Elnatan, P. Jaishankar, P. Bisignano, F.R. Garcia-Gonzalo, A.L. Krup, M. Eberl, E.F.X. Byrne, C. Siebold, S.Y. Wong, A.R. Renslo, M. Grabe, J.G. McDonald, L. Xu, P.A. Beachy, and J.F. Reiter. 2018. Cilia-Associated Oxysterols Activate Smoothened. Mol Cell. 72:316–327 e315.

Riobo, N.A., B. Saucy, C. Dilizio, and D.R. Manning. 2006. Activation of heterotrimeric G proteins by Smoothened. Proc Natl Acad Sci USA. 103:12607–12612.

Santos, N., and J.F. Reiter. 2014. A central region of Gli2 regulates its localization to the primary cilium and transcriptional activity. J Cell Sci. 127:1500–1510.

Sarkisian, M.R., D. Siebzehnrubl, L. Hoang-Minh, L. Deleyrolle, D.J. Silver, F.A. Siebzehnrubl, S.M. Guadiana, G. Srivinasan, S. Semple-Rowland, J.K. Harrison, D.A. Steindler, and B.A. Reynolds. 2014. Detection of primary cilia in human glioblastoma. J Neurooncol. 117:15–24.

Shao, J., L. Xu, L. Chen, Q. Lu, X. Xie, W. Shi, H. Xiong, C. Shi, X. Huang, J. Mei, H. Rao, H. Lu, N. Lu, and S. Luo. 2017. Arl13b promotes gastric tumorigenesis by regulating Smo trafficking and activation of the hedgehog signaling pathway. Cancer Res. 77:4000–4013.

Shen, F., L. Cheng, A.E. Douglas, N.A. Riobo, and D.R. Manning. 2013. Smoothened is a fully competent activator of the heterotrimeric G protein G(i). Mol Pharmacol. 83:691–697.

Shi, P., L.B. Hoang-Minh, J. Tian, A. Cheng, R. Basrai, N. Kalaria, J.J. Lebowitz, H. Khoshbouei, L.P. Deleyrolle, and M.R. Sarkisian. 2021. HDAC6 Signaling at Primary Cilia Promotes Proliferation and Restricts Differentiation of Glioma Cells. Cancers (Basel). 13:1644.

Shi, P., J. Tian, B.S. Ulm, J.C. Mallinger, H. Khoshbouei, L.P. Deleyrolle, and M.R. Sarkisian. 2022. Tumor Treating Fields Suppression of Ciliogenesis Enhances Temozolomide Toxicity. Front Oncol. 12:837589.

Shireman, J.M., F. Atashi, G. Lee, E.S. Ali, M.R. Saathoff, C.H. Park, S. Savchuk, S. Baisiwala, J. Miska, M.S. Lesniak, C.D. James, R. Stupp, P. Kumthekar, C.M. Horbinski, I. Ben-Sahra, and A.U. Ahmed. 2021. De novo purine biosynthesis is a major driver of chemoresistance in glioblastoma. Brain. 144:1230–1246.

Stupp, R., W.P. Mason, M.J. van den Bent, M. Weller, B. Fisher, M.J. Taphoorn, K. Belanger, A.A. Brandes, C. Marosi, U. Bogdahn, J. Curschmann, R.C. Janzer, S.K. Ludwin, T. Gorlia, A. Allgeier, D. Lacombe, J.G. Cairncross, E. Eisenhauer, and R.O. Mirimanoff. 2005. Radiotherapy plus concomitant and adjuvant temozolomide for glioblastoma. N Engl J Med. 352:987–996.

Tykocki, T., and M. Eltayeb. 2018. Ten-year survival in glioblastoma. A systematic review. J Clin Neurosci. 54:7–13.

Wang, Y., Z. Zhou, C.T. Walsh, and A.P. McMahon. 2009. Selective translocation of intracellular Smoothened to the primary cilium in response to Hedgehog pathway modulation. Proc Natl Acad Sci USA. 106:2623–2628.

Wei, L., W. Ma, H. Cai, S.P. Peng, H.B. Tian, J.F. Wang, L. Gao, and J.P. He. 2022. Inhibition of Ciliogenesis Enhances the Cellular Sensitivity to Temozolomide and Ionizing Radiation in Human Glioblastoma Cells. Biomed Environ Sci. 35:419–436.

Wheway, G., L. Nazlamova, and J.T. Hancock. 2018. Signaling through the Primary Cilium. Front Cell Dev Biol. 6:8.

Xu, Q., X. Yuan, G. Liu, K.L. Black, and J.S. Yu. 2008. Hedgehog signaling regulates brain tumor-initiating cell proliferation and portends shorter survival for patients with PTEN-coexpressing glioblastomas. Stem Cells. 26:3018–3026.

Yang, W., Y. Liu, R. Gao, H. Yu, and T. Sun. 2018. HDAC6 inhibition induces glioma stem cells differentiation and enhances cellular radiation sensitivity through the SHH/Gli1 signaling pathway. Cancer Lett. 415:164–176.

Ye, F., A.R. Nager, and M.V. Nachury. 2018. BBSome trains remove activated GPCRs from cilia by enabling passage through the transition zone. J Cell Biol. 217:1847–1868.

Zalenski, A.A., S. Majumder, K. De, and M. Venere. 2020. An interphase pool of KIF11 localizes at the basal bodies of primary cilia and a reduction in KIF11 expression alters cilia dynamics. Sci Rep. 10:13946.

